# Decoupling from yolk sac is required for extraembryonic tissue spreading in the scuttle fly *Megaselia abdita*

**DOI:** 10.1101/236364

**Authors:** Francesca Caroti, Everardo González Avalos, Viola Noeske, Paula González Avalos, Dimitri Kromm, Maike Wosch, Lucas Schütz, Lars Hufnagel, Steffen Lemke

## Abstract

Extraembryonic tissues contribute to animal development, which often entails spreading over embryo or yolk. Apart from changes in cell shape, the requirements for this tissue spreading are not well understood. Here we analyze spreading of the extraembryonic serosa in the scuttle fly *Megaselia abdita*. The serosa forms from a columnar blastoderm anlage, becomes a squamous epithelium, and eventually spreads over the embryo proper. We describe the dynamics of this process in long-term, whole-embryo time-lapse recordings, demonstrating that free serosa spreading is preceded by a prolonged pause in tissue expansion. Closer examination of this pause reveals mechanical coupling to the underlying yolk sac, which is later released. We find mechanical coupling prolonged and serosa spreading impaired after knockdown of *M. abdita Matrix metalloprotease 1*. We conclude that tissue-tissue interactions provide a critical functional element to constrain spreading epithelia.

**Impact Statement:** Extraembryonic tissue spreading in the scuttle fly *Megaselia abdita* requires mechanical decoupling from the underlying yolk sac.

## INTRODUCTION

In the early stages of animal development, pre-patterned cells are collectively set aside from the embryo proper to differentiate into specialized, extraembryonic epithelia, which then generate a local environment and support the growing organism from outside the embryo (Wolpert and Tickle, 2011). In chick, quails, frogs, fish and other vertebrates with a yolk-rich egg, such extraembryonic epithelia typically expand from the periphery of the embryo, spread over, and eventually envelope the underlying yolk cell (Downie, 1976; Futterman et al., 2011; Keller, 1980; Trinkaus, 1951; Arendt and Nübler-Jung, 1999). In bugs, beetles, butterflies, mosquitoes and other insects, extraembryonic epithelia already develop on top of a yolk sac, and when they spread, they eventually envelope the embryo proper (Panfilio et al., 2006; Handel et al., 2000; Kraft and Jäckle, 1994; Goltsev et al., 2009; Anderson, 1972a; Anderson, 1972b). To form such epithelial envelopes, extraembryonic tissues dramatically expand their area and often undergo a transformation from a columnar architecture with thin and tall cells to a squamous epithelium with spread-out and short cells (Anderson, 1972a; Anderson, 1972b; Bruce, 2016).

While synchronized and cell-autonomous flattening of cells is likely a necessary component, work from the fruit fly *Drosophila melanogaster* suggests that cell shape changes alone may not be sufficient to explain extraembryonic tissue spreading in insects (Lacy and Hutson, 2016). In *D. melanogaster*, cells of the extraembryonic amnioserosa synchronously and autonomously change their shape and thereby transform an initially columnar tissue into a squamous epithelium (Pope and Harris, 2008; Campos-Ortega and Hartenstein, 1997). In contrast to the extraembryonic tissues in most other insects, however, the amnioserosa does not extend from its dorsal position over the yolk and never spreads over the embryo (Panfilio, 2008; Schmidt-Ott and Kwan, 2016). These observations suggest that additional cellular mechanisms are required to instruct or permit extraembryonic tissue spreading.

To identify unknown mechanisms required for extraembryonic tissue spreading in insects, we investigated early development in the scuttle fly *Megaselia abdita*. *M. abdita* shared a last common ancestor with *D. melanogaster* about 150 million years ago (Wiegmann et al., 2011). While overall embryonic development of *M. abdita* and *D. melanogaster* is conserved and comparable (Wotton et al., 2014), the two species differ in their extraembryonic development (Rafiqi et al., 2008; Wotton et al., 2014). In *D. melanogaster*, the amnioserosa is set up as a single extraembryonic epithelium by expression of the Hox3 transcription factor Zerknüllt (Zen), which is controlled by peak levels of BMP signaling along the dorsal midline of the blastoderm embryo (Gavin-Smyth and Ferguson, 2014). In contrast to *D. melanogaster, M. abdita* develops two distinct extraembryonic tissues, the serosa, and bordering it, the amnion (Kwan et al., 2016; Rafiqi et al., 2008; Rafiqi et al., 2012). In the *M. abdita* blastoderm embryo, expression of the *zen* orthologue defines the serosa anlage along the dorsal midline (Rafiqi et al., 2008). Adjacent to this *zen*-defined serosa anlage, but within the range of dorsal BMP signaling, a narrow domain void of both *zen* and general embryonic patterning genes was identified as putative amnion anlage (Figure 1A; (Kwan et al., 2016; Rafiqi et al., 2012)). Cells of the serosa and amnion anlage undergo synchronous cell shape changes and eventually differentiate into squamous epithelia. The serosa then separates from the adjacent amnion, spreads freely, and continuously increases its cell size until it eventually closes as perfect envelope on the ventral side of the egg (Rafiqi et al., 2008; Rafiqi et al., 2010). Because *M. abdita* has retained critical elements of ancestral extraembryonic tissue spreading, it has been previously identified as key organism to understand the evolutionary origin of the amnioserosa as a single, non-spreading extraembryonic epithelium (Hallgrímsson et al., 2012).

**Figure 1.**
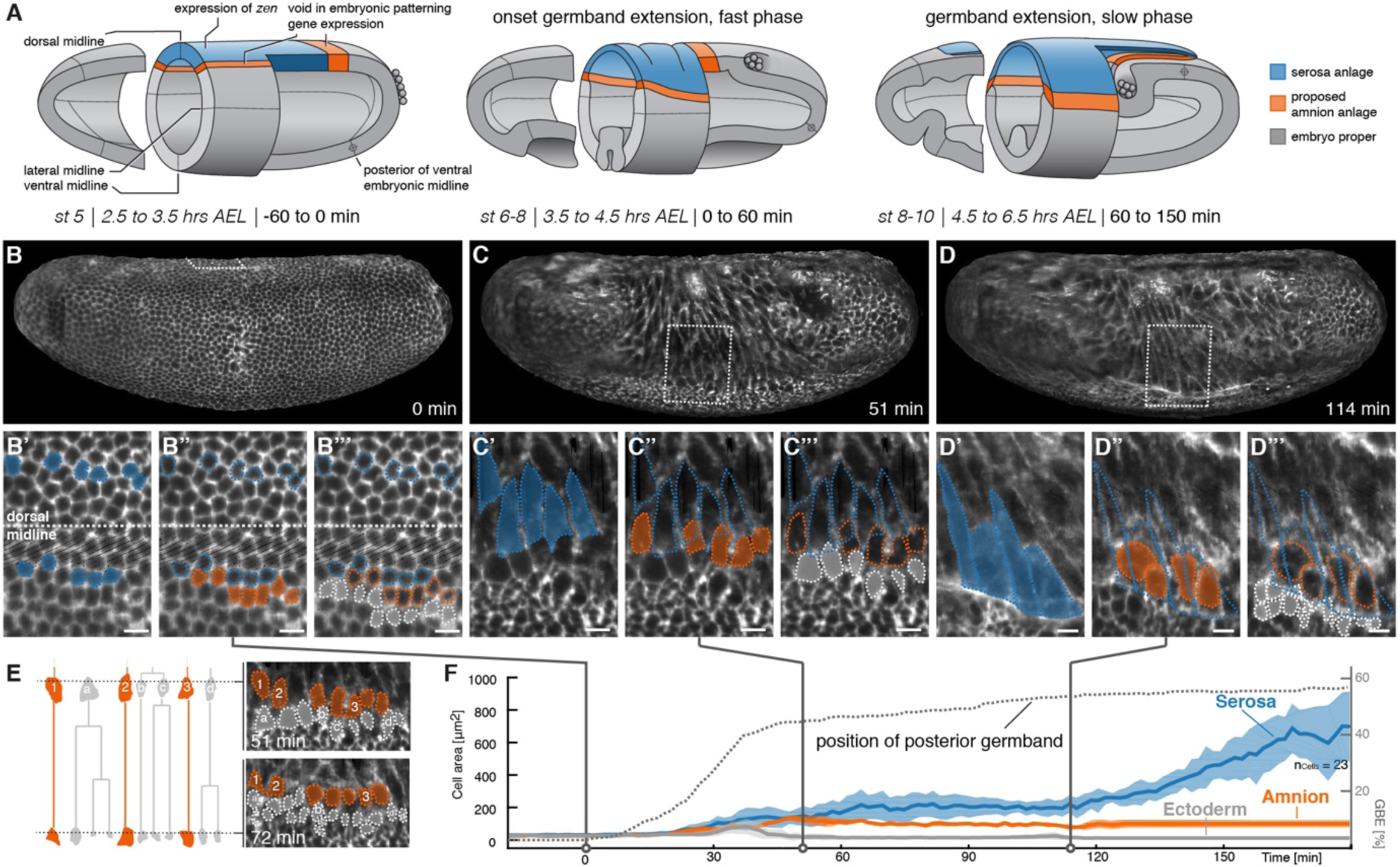
Tracking of blastoderm cells characterizes serosa and amnion differentiation. (**A**) Model of early extraembryonic tissue development in *M. abdita* based on marker gene expression in fixed specimen. Stage (st) and time after egg lay (AEL) are defined as in (Wotton et al., 2014); absolute time given in minutes relative to the onset of germband extension (onset GBE = 0 min). (**B-D’’’**) Global embryonic views of SPIM recorded embryos at corresponding stages (**B-D**), with tracked and marked serosa (blue), amnion (orange), and ectoderm cells (grey) in 2D-projections of indicated surface areas in dorsal (**B’-B’’’**) and lateral views (**C’-C’’’, D’-D’’’**). Cells of serosa were identified based on their ability to spread over the embryo and then tracked back to the cellular blastoderm. (**E**) Cell lineage and divisions in putative amnion and ectoderm cells. Cells directly adjacent to the serosa never divided and could be back-tracked to a single row of cells next to the serosa anlage; these cells were classified as presumptive amnion. Cells further distal to the serosa divided, eventually decreased in cell size, and were classified as presumptive ectoderm. (**F**) Quantitative analysis of cell size of tracked serosa, amnion, and ectodermal cells relative to GBE as measure of developmental progression. The position of the posterior germband is indicated in % egg length (0% = posterior pole; dotted line); Standard error of mean shown as shades. Unless indicated otherwise, embryos and close-ups are shown with anterior left and dorsal up. Scale bars, 10 µm.

To take advantage of its close relationship to *D. melanogaster* and use it as model to address cell-biological mechanisms of extraembryonic tissue spreading, tissue and cell dynamics in *M. abdita* development are required to be studied with high spatiotemporal resolution. Here we have established time-lapse recordings at the necessary resolution in injected embryos using confocal and light sheet microscopy. This has allowed us to address open questions regarding amnion and serosa development, and it identified mechanical coupling between serosa and yolk sac as a critical element to control serosa spreading. We genetically interfered with serosa coupling in *M. abdita* and found that changes in tissue-tissue interaction provide a compelling variable for the evolution of epithelial spreading.

## RESULTS

### ***In toto*** time-lapse **recordings faithfully recover known landmarks of embryonic and extraembryonic development in *M. abdita***

Extraembryonic development in *M. abdita* has been characterized by the formation of two extraembryonic tissues, the amnion and the serosa. Both tissues develop from columnar cells of the blastoderm embryo and have been associated with dramatic changes in cell size. To test whether such cellular properties could be used to trace amnion and serosa differentiation from the blastoderm stage onwards, we carefully examined fixed specimen by sampling subsequent time points of development using precisely staged depositions (Figure 1 supplement 1). Our quantitative analyses of cell shapes clearly showed two classes of cells, i.e. large and flat and presumably extraembryonic cells, and small and round and presumably ectodermal cells (Figure 1 supplement 1). However, our data was not sufficient to distinguish between putative amnion and serosa cells, which made it impossible to follow their development from the blastoderm embryo.

Time lapse recordings of individual living specimen permit the tracing of cells and tissues throughout development (Keller, 2013), which avoids the discontinuity that limited our analysis of fixed specimen. In case of extraembryonic development in *M. abdita*, this approach would require imaging of the entire embryo (the serosa is specified on the dorsal side and eventually fuses on the ventral side (Rafiqi et al., 2008)), at cellular resolution, and for an extended period of time. Time lapse recordings of embryonic development using selective plane illumination microscopy (SPIM) have been previously shown to provide the spatiotemporal resolution necessary for similar analyses (Chhetri et al., 2015; Rauzi et al., 2015; Wolff et al., 2018), suggesting that SPIM could also be used to explore formation and differentiation of extraembryonic tissues in the *M. abdita* embryo.

To establish SPIM imaging for *M. abdita* embryos, we first established fluorescent markers for cell outline and nuclei (Lifeact and Histone H1, see material and methods). Next, we tested whether overall development was affected by long-term imaging of injected *M. abdita* embryos. To assess the putative impact of extended time-lapse recordings, we compared universal landmarks of fly development (onset of germband extension, onset of germband retraction, and end of germband retraction) in SPIM recordings with our own live observations as well as previously described recordings of *M. abdita* development (Wotton et al., 2014). For all three landmarks, our live observations confirmed previously described timing (onset of germband extension 3:45 hrs after egg laying (AEL), onset of germband retraction 8:10 hrs AEL, and end of germband retraction 11:05 hrs AEL (Wotton et al., 2014)) within a window of 15 min (n = 24), which corresponded to the time interval of egg deposition. In our quantified SPIM recordings we found the same timing of events, suggesting that development of injected *M. abdita* embryos was not notably affected by long-term SPIM imaging. In the following, we will use previously defined staging and terminology for *M. abdita* development where possible (Rafiqi et al., 2008; Wotton et al., 2014), while absolute times will be provided relative to the onset of germband extension (0 min).

To follow development and differentiation of *M. abdita* serosa and amnion from their proposed blastoderm anlage, we selected a representative time-lapse recording for cell tracking and followed cells of putative serosa, amnion, and ectoderm throughout early development (Figure 1B-D’’’). The serosa developed from a dorsal anlage that was about six to seven cells wide (Figure 1B,B’). These cells did not divide, increased in cell surface area (Figure 1C,C’), and eventually spread over the adjacent tissue (Figure 1D,D’). The width of the inferred serosa domain perfectly coincided with gene expression of the known serosa key regulator, the homeodomain transcription factor Zerknüllt (Zen), while the observed cell dynamics matched serosa behavior as previously inferred from fixed specimen (Rafiqi et al., 2008). Directly adjacent to the unambiguously identified serosa anlage, we identified a single row of cells (Figure 1B,B’’), which increased in apical cell area (Figure 1C,C’’) but eventually did not become part of the spreading serosa (Figure 1D,D’’). These cells did not divide (Figure 1E), and, consistent with patterning information in the early blastoderm embryo (Figure 1A; (Kwan et al., 2016; Rafiqi et al., 2012)), we propose that they formed the presumptive amnion. Next to this putative amnion, cells increased in apical cell area temporarily (Figure 1B,B’’’,C,C’’’) but eventually divided (Figure 1D,D’’’,E) and decreased in size, suggesting differentiation into ectodermal cells. The overall analyses of surface cell area at later stages of development, i.e. after germband extension, indicated a substantial increase in surface area for serosa cells (up to 10-fold), a still notable increase for presumptive amnion cells (about 2-fold), and a slight reduction of cell surface area in the ectoderm (Figure 1F). Taken together, our cell tracking analyses support the previously proposed model of the *M. abdita* blastoderm fate map and set the stage for a more global analysis of amnion and serosa tissue behavior.

### The amnion in *M. abdita* develops as a lateral stripe of cells

Previous analyses of *M. abdita* amnion development have led to conflicting hypotheses regarding its position and topology. Based on either cell outline or marker gene expression, the *M. abdita* amnion has been suggested to consist of either a dorsally closed epithelium or a thin lateral stripe of cells (Kwan et al., 2016; Rafiqi et al., 2008). To resolve this question, we aimed to follow expansion and development of the presumptive amnion in our time-lapse recordings. To achieve this in the absence of a specific *in vivo* reporter, we developed a set of image processing routines for embryos recorded in isotropic 3D volumes. These routines allowed us first to identify the surface of the embryo and to digitally "peel off" individual layers until the serosa was removed. We then flattened the surface into a two dimensional carpet and marked the remaining cells with large surface area as presumptive amnion. Finally, these carpets were projected again into their initial three dimensional shape to provide three dimensional renderings (Figure 2 supplement 1; Material and Methods).

Thus following development of the presumptive amnion through consecutive stages of development, our results suggested that the *M. abdita* amnion consisted of a lateral tissue, which was essentially one cell wide, and a cap at the posterior end of the germband that closed over the ventral side of the embryo (which, because of the extended germband, faced the dorsal side of the egg, Figure 2A-D). During germband extension and up until the onset of germband retraction, cells of the presumptive lateral amnion had a rather smooth outline and appeared to be folded over the ectoderm; with the onset of germband retraction, these cells developed notable protrusions and appeared to crawl over the yolk sac (Figure 2C,D). To follow the position of the presumptive amnion more precisely, we tracked individual amnion cells along the embryonic circumference over the course of germband extension and retraction. Our analyses further consolidated the notion of a lateroventral amnion in *M. abdita* (Figure 2E,F).

**Figure 2.**
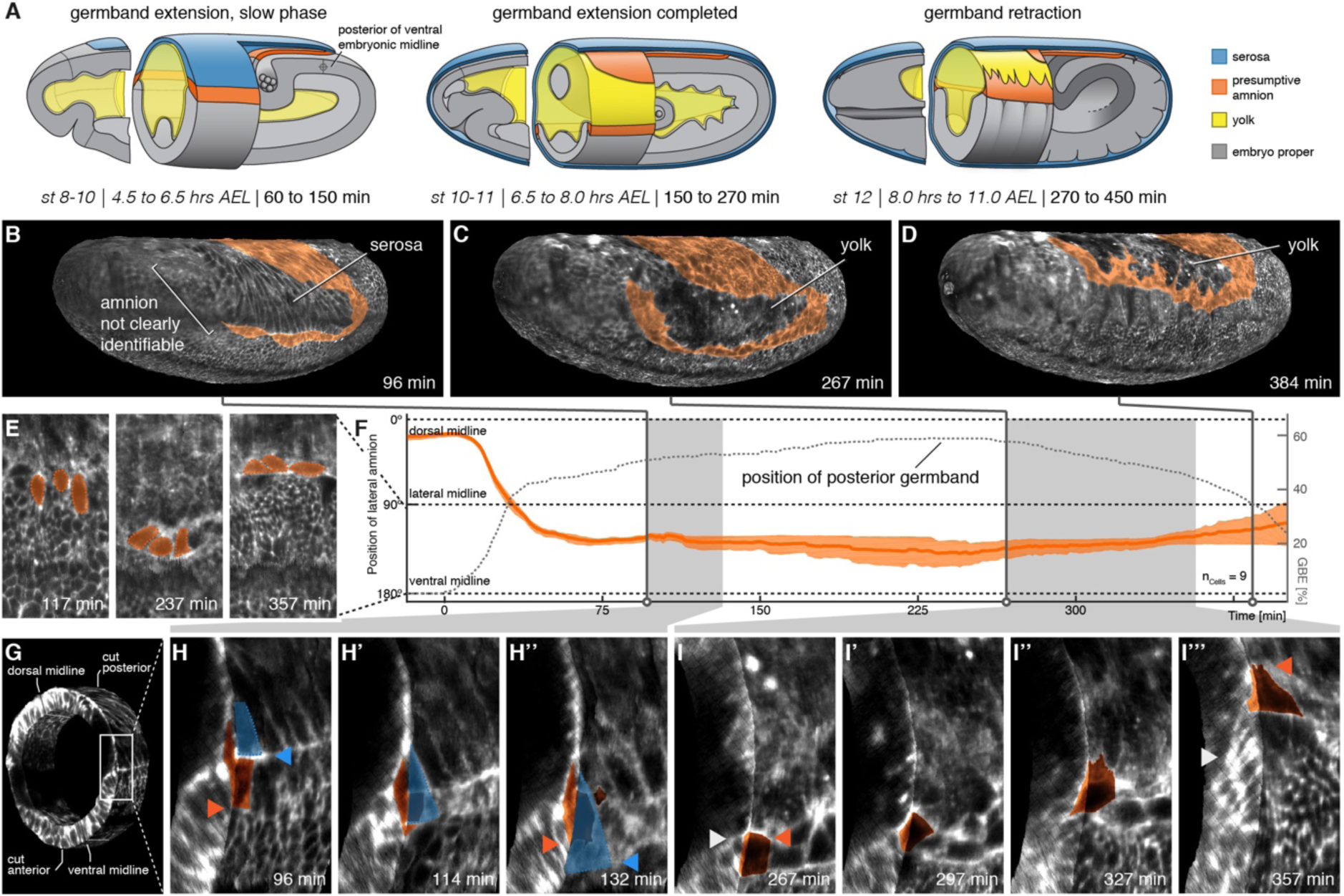
The lateral anlage of the *M. abdita* putative amnion differentiates into a one-to-two cell wide lateral epithelium. (**A**) Model of extraembryonic tissue development in *M. abdita* based on SPIM time-lapse recordings; staging as in Figure 1. (**B-D**) Global embryonic views of SPIM recorded embryos at corresponding stages. To reveal and mark the putative amnion, the surface layer has been digitally removed (Figure 2 supplement 1). (**E,F**) Tracking (**E**) and plotting (**F**) of amnion cell position (orange) along the dorso-ventral circumference (0° corresponds to dorsal, 180° to ventral midline) relative to GBE as measure of developmental progression (dotted line), standard error of mean shown as shades. (**G-I’’’**) Donut section of SPIM recorded image volume (**G**) to illustrate close-up views of amnion cell behavior as the serosa (blue) detaches during germband extension (**H**) and after onset of germband retraction (**I**). As the serosa spreads over the amnion (**H-H’’**), the amnion appears to bend underneath the serosa. The serosa then separates from the amnion and spreads over the embryo proper. During germband retraction (**I-I’’’**), the amnion starts to extend actin-rich protrusions and leads the ectoderm as the tissue progresses towards the dorsal midline. Triangles indicate relative positions of amnion, serosa, and ectoderm in first and last time points (amnion = orange, serosa = blue, and embryonic cells = grey). Embryos are shown with anterior left and dorsal up.

To understand how amnion cells reacted after disjunction from the serosa, we computed donut-like sections of the developing embryo that allowed us to observe embryo surface and transverse section in 3D renderings (Figure 2G). In these renderings, the serosa first separated from the presumptive amnion and then appeared to drag it slightly over the adjacent ectoderm. As a result, amnion cells seemed to be turned with their lateral side towards the basal membrane of the serosa (Figure 2H). This orientation was reversed during germband retraction; presumptive amnion cells no longer appeared folded over the adjacent ectoderm and instead showed a crawling-like behavior with long cellular protrusions over the yolk sac (Figure 2D,H).

### The serosa in *M. abdita* expands in distinct phases

Previous analyses of *M. abdita* serosa expansion have been based on the expression of specific marker genes in fixed embryos (Rafiqi et al., 2008) as well as on bright field microscopy time-lapse recordings (Wotton et al., 2014). These analyses suggest that the serosa starts to be detectable about 30-45 min after the onset of germband extension; it then expands, presumably homogeneously, for almost 2.5 hrs until the tissue front passes the equator at both poles of the embryo; and then, within minutes, it fuses rapidly along the ventral midline (Wotton et al., 2014).

We revisited these dynamics in our SPIM recordings by marking the serosa in projected carpets, similar as outlined above for the amnion. Marking of the serosa was guided by cell size as well as the enrichment of the actin reporter at the interface of amnion and serosa (Figure 4 supplement 1), which indicated the formation of a supracellular actin cable at the serosa boundary similar to the actin cable observed during serosa formation in *T. castaneum* (Benton and St Johnston, 2003). Following marking of the serosa, we identified first signs of tissue expansion after about 35 min (Figure 3A). The serosa approached the equator at the poles at about 150 min (Figure 3B) and passed it 15 min later; ventral fusion along the midline was then completed after 240 min (Figure 3C). Our measures of serosa area increase were overall in accordance with previous analyses (Figure 3D,E). The increased resolution in ventral views allowed us to determine the specific dynamics of ventral fusion and suggested a closure rate of about 11 µm/min along the anterior-to-posterior axis of the serosal window, and 3 µm/min across (Figure 3F,G), which was considerably faster than rates that have been described for dorsal closure of the amnioserosa in *D. melanogaster* (Hutson et al., 2003). Notably, our continuous mapping of area expansion indicated that *M. abdita* serosa spreading was a non-homogeneous process, in which two periods of almost linear area growth were interrupted by a roughly 50-minute interval without substantial tissue area increase (70-120 min, Figure 3D). At the end of this interval, we observed disjunction of the serosa from the embryo proper, first at the anterior, then at the posterior and finally at its lateral sides (Figure 3E).

**Figure 3.**
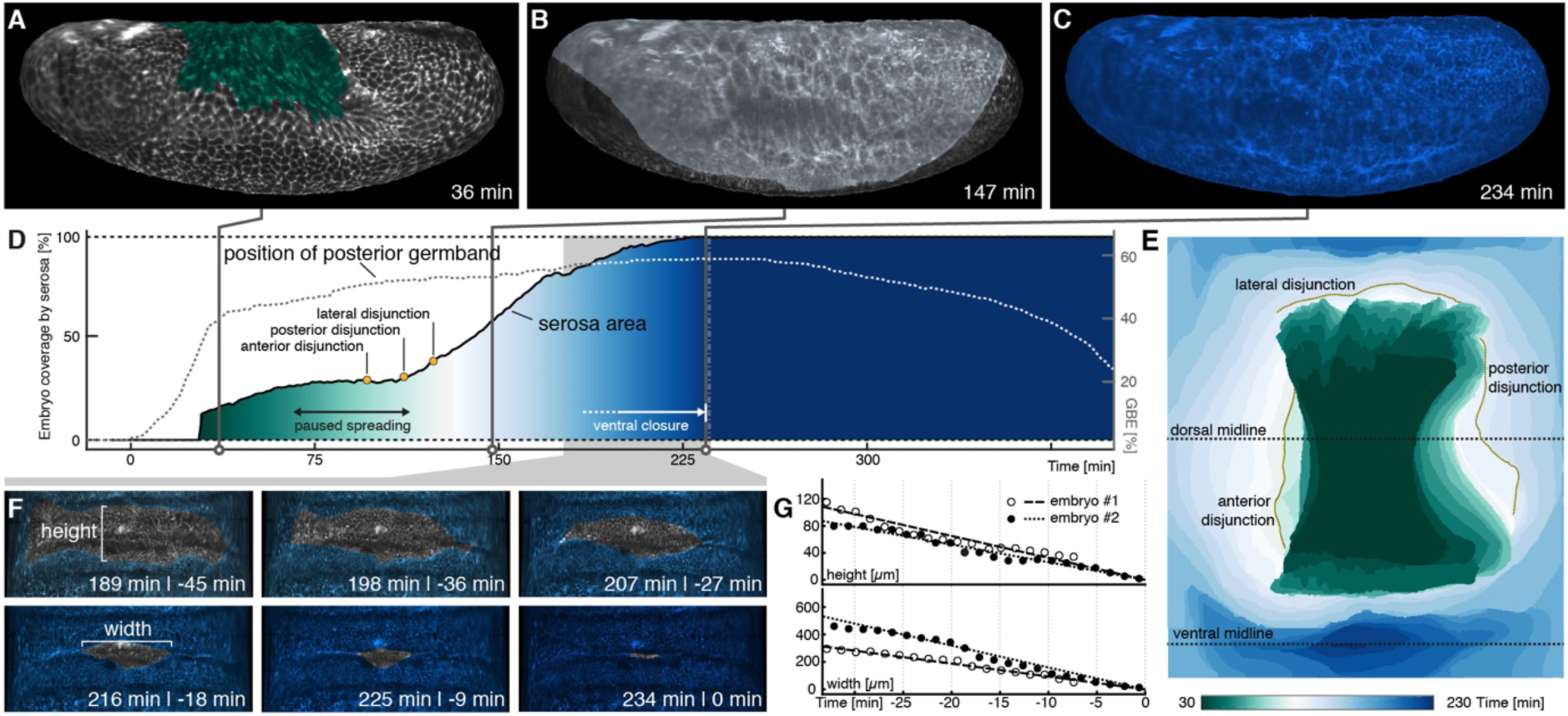
Serosa expansion in *M. abdita* is characterized by two distinct phases that are separated by a pause in tissue spreading. (**A-C**) Onset (**A**), passing of poles (**B**), and ventral closure (**C**) of marked serosa illustrate previously described stages of expansion (Wotton et al., 2014). (**D,E**) Plotting of serosa area over time and relative to germband extension (**D**) indicates two additional stages of serosa expansion, i.e. a pause in spreading and the subsequent disjunction, first at the anterior, then posterior, and finally lateral circumference (**E**). Time is indicated in colormap. (**F,G**) Progression of ventral closure indicated in projections of ventral embryo surfaces (0 = ventral closure) (**F**) and quantified as linear decrease in height and width of serosal window (**G**).

### De-coupling of serosa and yolk sac is preceding serosa tissue spreading

The observed interruption in serosa area increase hinted towards a possible mechanism of interfering with tissue spreading. For example, the interruption in serosa expansion could be explained by a temporal pause in the tissue-autonomous program of cell thinning and spreading. However, expression of the tissue-fate determining Hox-3 transcription factor Zen remains high also throughout stages in which tissue expansion paused (Rafiqi et al., 2008), indicating little, if any, change in the upstream genetic regulation of cell thinning and spreading. Alternatively, serosa area increase could be paused if further tissue spreading was hindered by adhesion to an underlying substrate. A possible candidate for such a substrate is the yolk sac.

To test whether interaction between yolk sac and serosa could explain the pause in tissue spreading, we aimed to test for mechanical interaction between yolk sac and serosa. To be able to visualize the yolk sac, we first cloned the *M. abdita* orthologue of *basigin* (*Mab-bsg*, Figure 4 supplement 1), which encodes a trans-membrane protein that in *D. melanogaster* is enriched in the yolk sac membrane (Reed et al., 2014; Goodwin et al., 2016). Analysis of *Mab-bsg* expression indicated that the gene was expressed predominantly in the periphery of the yolk sac (Figure 4 supplement 1). We then asked whether a fluorescent Basigin reporter (Basigin-eGFP) could be used to visualize the yolk sac membrane like in *D. melanogaster* (Goodwin et al., 2016). To analyze expression of Basigin-eGFP together with ubiquitously marked membranes, we first injected Lifeact-mCherry into early syncytial embryos. Then, to restrict Basigin-eGFP to the yolk sac, we injected capped mRNA encoding the fusion protein after germband extension had started and cellularization was presumably complete (Rafiqi et al., 2010). Visual inspection of time-lapse recordings indicated that fluorescent signals in serosa and yolk sac could be separated, allowing us to distinguish between movements in either tissue (Figure 4A-C).

**Figure 4.**
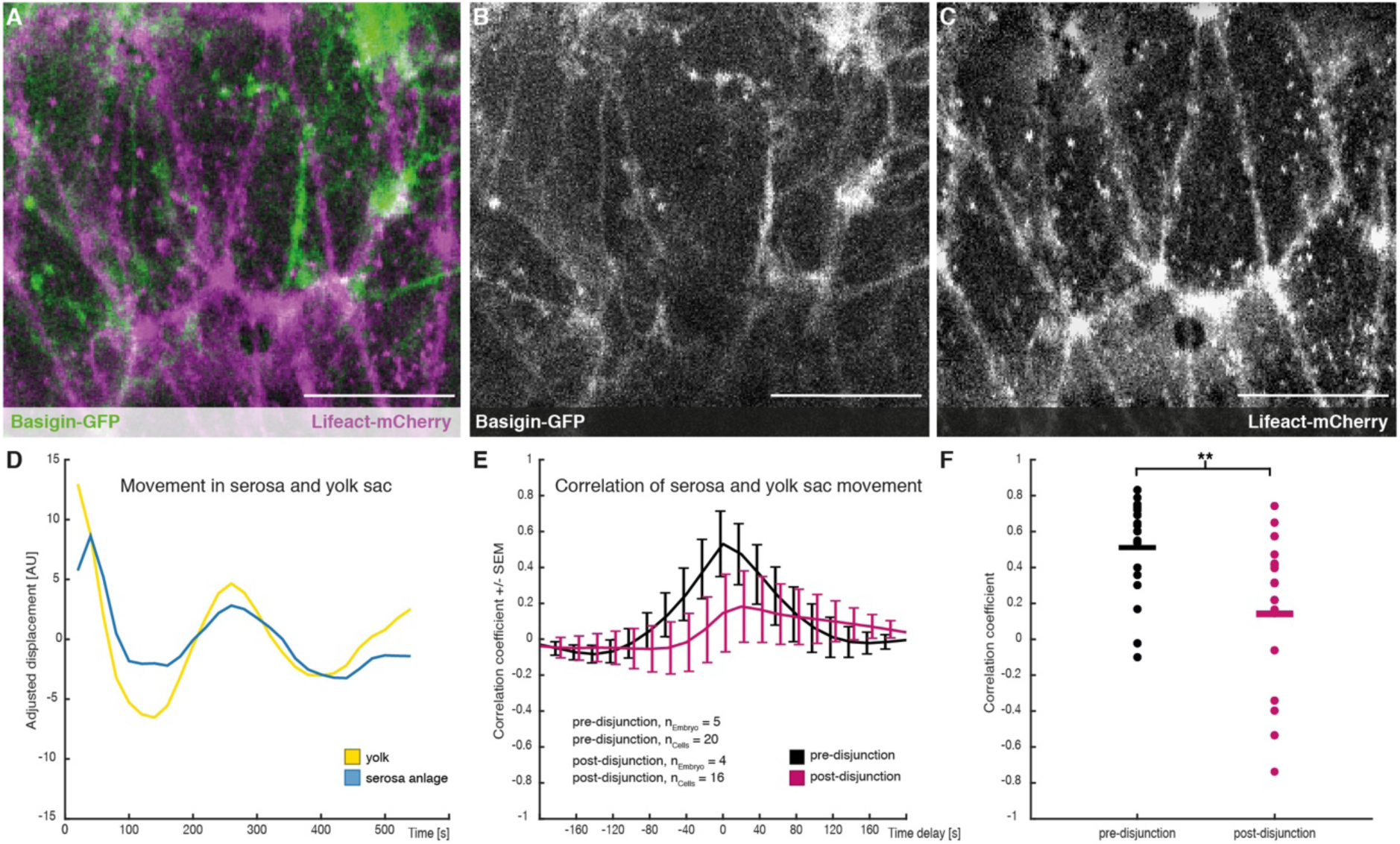
Cross-correlation of serosa cell and yolk sac movements suggest a decoupling of serosa prior to free spreading. (**A-C**) Images of serosa (visualized with Lifeact-mCherry) and underlying yolk sac (expressing Basigin-eGFP) are shown in average-intensity projections. (**D**) Adjusted serosa (blue) and yolk sac (yellow) displacement measured by optical flow analysis exemplary for one serosa cell and corresponding substrate area underneath. (**E)** Average cross-correlation of cell and substrate movements indicate coupling before (black) and decoupling after (magenta) disjunction of the serosa from the embryo proper. Standard error of mean shown as bars. (**F**) Collective comparison of correlation coefficients for individual cells before and after disjunction of the serosa from the embryo, bar indicates the mean. ** p = 0.00686 based on students t-test. Scale bars, 20 µm.

With Basigin-eGFP established as fluorescent reporter for the yolk sac membrane in *M. abdita*, we first asked whether this reporter could be used to detect movements at the yolk sac surface. To quantify these movements we measured the mean integrated flow velocity of Basigin in the yolk sac membrane using optical flow (see Material and Methods). These analyses detected presumably oscillatory membrane behavior along the anterior-to-posterior axis, which seemed to coincide with similar movements in individual serosa cells (Figure 4D). The direct comparison of movements in yolk sac and serosa revealed a strong positive correlation, which was very specific to individual serosa cells and the yolk sac membrane directly underneath (Figure 4E,F and Figure 4 supplement 1). Our findings indicated mechanical coupling of the two membranes, which we found significantly reduced after the second phase of serosa extension had started (Figure 4E,F). Taken together, our results suggest that free spreading of the *M. abdita* serosa over the embryo is preceded by a decoupling of the serosa from the underlying yolk sac.

### *Mab-Mmp1* modulates tissue-tissue interaction between yolk sac and serosa

In *D. melanogaster*, modulation of tissue-tissue adhesion as well as complete tissue-tissue decoupling has been previously associated with *Matrix metalloprotease 1* (*Mmp1*) activity (Diaz-de-la-Loza et al., 2018; Glasheen et al., 2010; LaFever et al., 2017; Srivastava et al., 2007). To test whether activity of this matrix metalloprotease was involved in mechanical decoupling of serosa and yolk sac, we cloned the *M. abdita* orthologue of *Mmp1* (*Mab-Mmp1*) and found it expressed in yolk sac nuclei (Figure 5A). To test whether *Mab-Mmp1* had an effect on interaction between serosa and yolk sac, we analyzed embryos in which gene activity was knocked down using RNAi or CRISPR. We first tested for putative effects on serosa development by staining embryos for the serosa specific marker gene *dopa decarboxylase* (*Mab-ddc*; (Rafiqi et al., 2010)). In wildtype embryos, the serosa closes about 7 to 7.5 hrs after egg lay (Figure 3D), which was reflected in uniform staining of *Mab-ddc* in correspondingly staged and fixed embryos (Figure 5B, (Rafiqi et al., 2010)). Following knockdown by RNAi (96%, n=100/104) and CRISPR (56%, n=77/136), *Mab-ddc* expression was reduced at corresponding stages to a dorsal domain, suggesting that serosa spreading was impaired (Figure 5C,D).

**Figure 5.**
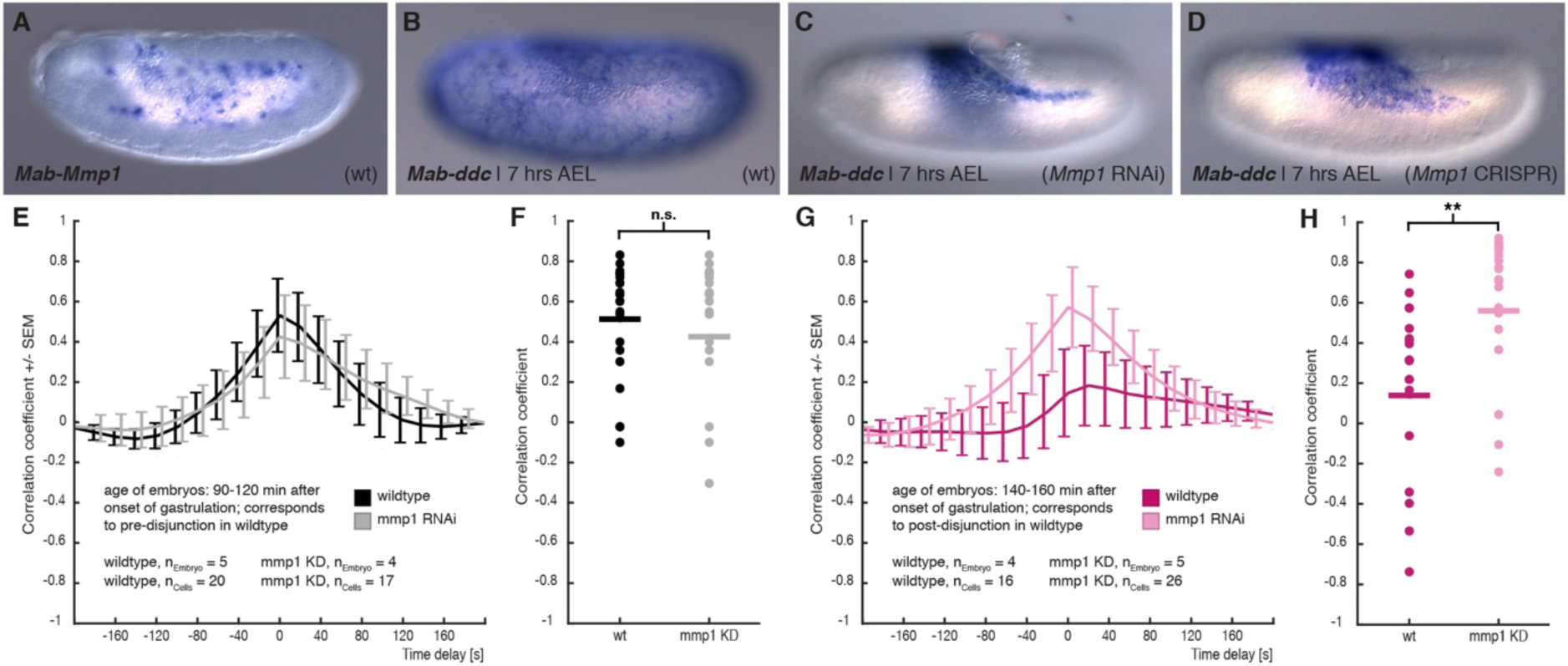
*M. abdita matrix metalloprotease 1* (*Mab-Mmp1*) regulates serosa decoupling. (**A**) Expression of *Mab-Mmp1* during germband extension stage. (**B-D**) Expression of *M. abdita dopa decarboxylase* (*Mab-ddc*) as serosa marker during germband extension stage in wildtype (**B**), in *Mab-Mmp1* RNAi (**C**), and *Mab-Mmp1* CRISPR/Cas9 embryos (**D**). (**E,F**) Average cross-correlation function of corresponding cell and substrate movements indicate similar level of mechanical coupling in wildtype (black) and *Mab-Mmp1* RNAi embryos (grey) at the end of serosa expansion pause in wildtype embryos (**E**), with statistical support from correlation coefficients of individual cells (**F**). (**G,H**) Average cross-correlation function of corresponding cell and substrate movements indicate low mechanical coupling in wildtype (dark magenta) and high mechanical coupling in *Mab-Mmp1* RNAi embryos (light magenta) after onset of free serosa expansion in wildtype embryos (**G**), with statistical support from correlation coefficients for individual cells (**H**). Bars indicate mean. ** p = 0.00279; n.s. p = 0.48746, based on student’s t-test.

To address how knockdown of *Mab-Mmp1* affected mechanical coupling of serosa and yolk sac, we quantified tissue-level oscillation and correlated flow velocity in RNAi embryos before and after presumptive decoupling. In *Mab-Mmp1* RNAi embryos corresponding to wildtype developmental stages prior to decoupling (90-120 min), the correlation of movements in serosa and yolk sac were comparable to wildtype and suggested mechanical coupling between the two membranes (Figure 5E,F). In *Mab-Mmp1* RNAi embryos corresponding to wildtype developmental stages after decoupling (140-200 min), correlation of serosa and yolk sac movements remained virtually unchanged and were significantly higher than in wildtype (Figure 5G,H). These results indicated that mechanical coupling between the two membranes was not broken up, thus explaining impaired serosa spreading.

To determine whether serosa spreading was attenuated, delayed, or completely halted after knockdown of *Mab-Mmp1*, we analyzed serosa spreading by marker gene expression in older embryos, i.e. 9 hrs and 11 hrs after egg lay (Figure 5 supplement 1). In 9-hour old *Mab-Mmp1* RNAi embryos, the serosa was not yet fused; in many embryos it expanded to the mid-lateral side (33%, n=27/83), and a few embryos had strong phenotype with a dorsal serosa (4%, n=3/83). In 11-hour old *Mab-Mmp1* RNAi embryos, the serosa was fused in most embryos (77%, n=30/39), while a few specimen still showed the non-fused phenotype (23%, n=9/39). These results suggested that serosa spreading was attenuated or delayed after knockdown of *Mab-Mmp1*. To explain the observed delay in the completion of serosa closure, we cannot exclude a possibly incomplete knockdown of *Mab-Mmp1* activity. However, we consider it more likely that *Mab-Mmp1* acts redundantly with yet unknown factors to release the serosa from its yolk sac attachment.

## DISCUSSION

With our work we established the scuttle fly *M. abdita* as functionally accessible model to dissect cellular dynamics of extraembryonic tissue spreading in flies. Imaging the early *M. abdita* embryo over an extended period of time has allowed us to place the dynamics of extraembryonic development into the context of the developing embryo, at a spatiotemporal resolution that is comparable to previous work in *D. melanogaster* and the flour beetle *T. castaneum* (Benton et al., 2013; Chhetri et al., 2015; Goodwin et al., 2016; Hilbrant et al., 2016). This made it possible to connect previously studied pattern formation and signaling in the blastoderm with progression of cell and tissue differentiation (Kwan et al., 2016; Rafiqi et al., 2008; Rafiqi et al., 2012). The combination of long-term and *in toto* imaging enabled us to address open questions regarding amnion and serosa formation in *M. abdita*, and it allowed us to identify tissue-tissue interactions between yolk sac and serosa as a mechanism that controls extraembryonic tissue spreading in *M. abdita*.

### *M. abdita* forms an open lateral amnion

Our cell and tissue tracking in SPIM recordings of early *M. abdita* development were consistent with previously reported expression of a late amnion marker gene, *Mab-*egr. (Kwan et al., 2016). Our time-lapse recordings and previously published *Mab-egr* expression indicated that the *M. abdita* amnion developed as an open and mostly lateral tissue. The tissue topology of the amnion in *M. abdita* was thus markedly different than the closed and ventral amnion that has been reported for non-cyclorrhaphan flies and most other insects (Panfilio, 2008; Schmidt-Ott and Kwan, 2016). Connected to the lateral amnion, the amnion at the posterior end of the germband consisted of more cells and closed over the ventral midline of the embryo proper. The size of the differentiated amnion tissue area thus correlated with the amnion anlage in the *M. abdita* blastoderm, which was larger at the posterior than at the lateral anlage (Kwan et al., 2016; Rafiqi et al., 2012). While the posterior anlage seemingly provided enough cells to generate a ventrally closed amnion, the lateral line of one to two cells apparently was not sufficient to cover the area on the ventral side of the embryo. The implications of such a reduced lateral amnion on embryonic fly development in general are difficult to assess, in part because the precise function of the ventral amnion in insects is still unknown (Panfilio, 2008). We speculate, however, that differences in amnion topology (lateral vs. ventral) could be explained if insects with a ventral amnion had a larger amnion anlage, or if amnion cells could proliferate. Alternatively, a ventral amnion may be formed if the area that had to be covered was smaller, e.g. if the germband was more condensed or even participated in the initial ventral cavity as reported for *T. castaneum* (Benton, 2018).

### *M. abdita* serosa spreading proceeds in distinct phases

Area tracking of *M. abdita* serosa development identified two distinct phases of tissue spreading. These two phases corresponded to either the early, “tethered” or a late, “freed” state of the serosa and were interrupted by a notable pause in tissue area expansion. In the first phase of serosa spreading, the serosa was still continuous with amnion and ectoderm and thus part of a coherent epithelium. During this phase, the initially columnar blastoderm cells changed their shape and became squamous. In the second phase, the serosa broke free, cells increased further in their apical area, the tissue spread over the entire embryo proper, and eventually fused along the ventral midline. The dynamics of this process have been previously difficult to observe, presumably because the ventral serosa is difficult to discern in DIC images. Here we report that ventral closure of the serosa in *M. abdita* was substantially faster than dorsal closure of the *D. melanogaster* amnioserosa, and we propose that the comparatively fast fusion rate in the serosa can be explained by lack of segmental alignment and time consuming zippering as observed for dorsal closure in *D. melanogaster* (Jacinto et al., 2002).

### Free serosa spreading in *M. abdita* requires decoupling from yolk sac

Closer analysis of the transition from paused to free serosa spreading provided evidence for a change in tissue-tissue interaction between serosa and the underlying yolk sac. Strong correlation of movements in serosa and yolk sac membranes could be observed until the posterior of the extending germband reached about the middle of the embryo and indicated tight coupling between the membranes during paused serosa spreading. After that, free serosa spreading started and the correlation of movements between serosa and yolk sac was significant reduced, indicating that onset of free serosa spreading coincided with a decoupling from the yolk sac. We found that coupling persisted in *Mab-Mmp1* RNAi embryos substantially longer, and tissue-tissue interactions between yolk sac and serosa remained comparable to that of wildtype embryos during paused serosa spreading. Complementing these findings, we found serosa spreading impaired in *Mab-Mmp1* RNAi embryos. Taken together, our results strongly suggest that free serosa spreading in *M. abdita* requires its decoupling from the yolk sac.

Similar interactions between yolk sac and extraembryonic tissue have been reported previously in *D. melanogaster*, albeit at later stages of development, where they contribute to germband retraction and dorsal closure (Goodwin et al., 2016; Narasimha and Brown, 2004; Reed et al., 2004; Schöck and Perrimon, 2003). More generally, interactions of yolk sac and overlying epithelia have been long implicated in insect as well as vertebrate development (Anderson, 1972a; Anderson, 1972b; Bruce, 2016; Counce, 1961; Schmidt-Ott and Kwan, 2016), suggesting that yolk sac dependent regulation of serosa spreading in the *M. abdita* embryo may reflect a more common phenomenon.

### On the origin of the amnioserosa

Genetic changes that affect tissue contact remodeling for serosa and yolk sac in *M. abdita* could have similarly occurred in the last common ancestor of *D. melanogaster* and basal cyclorrhaphan flies with separate amnion and serosa. We speculate that a gradual loss of tissue decoupling between serosa and yolk sac could have resulted in impaired serosa spreading and thus restricted extraembryonic tissues to an exclusively dorsal position on top of the yolk sac. Topologically, and presumably also functionally, the resulting composite tissue of a non-spreading serosa and peripheral amnion would have shared similarities with the amnioserosa of *D. melanogaster*. Consistent with such a scenario, *D. melanogaster* lacks *Mmp1* expression in the yolk sac, and mutant embryos do not show a phenotype (Page-McCaw et al., 2003). Complementing biomechanical and gene expression analyses in the amnioserosa also support the idea that the amnioserosa has been, or still is, a composite tissue (Gorfinkiel et al., 2009; Wada et al., 2007). Against this background, our results illustrate how the *D. melanogaster* amnioserosa may have originated not by transformation of genetic patterning or tissue differentiation, but rather ancient changes in tissue-tissue interactions.

## MATERIAL AND METHODS

### Fly stocks

The laboratory culture of *Megaselia abdita* (Sander strain) was maintained at 25°C and a constant 16/8-hr day/night cycle as described previously (Caroti et al., 2015).

### Cloning and RNA synthesis

*Mab-Mmp1* was identified from genome and transcriptome sequences. A genomic fragment (GenBankID01) was cloned after PCR amplification from the locus, a cDNA fragment after amplification through 5’-RACE (GenBankID02). Double-stranded RNA (dsRNA) was synthesized as described (Urbansky et al., 2016); *Mab-Mmp1* dsRNA comprised pos. +103 to +1167 of the genomic fragment (pos. 1 refers to first nucleotide in ORF) and included a 57bp intron at pos. +575. Guide RNAs for a knock-out of *Mab-Mmp1* were designed using CCTop as CRISPR/Cas9 target online predictor (Stemmer et al., 2015). Three single guide RNAs (sgRNAs) were designed to target the following positions (pos. 1 refers to first nucleotide in ORF): sgRNA1, 5’-TGCAGAGCGTATCTCTTT, pos +404 to +387; sgRNA2, 5’-CGTGGACTATTGATTGTC, pos +710 to +693; sgRNA3, 5’-TCGGCAACCGAGTTTTCA, +898 to +881. Guide RNAs as well as Cas9 mRNA were synthesized as described (Stemmer et al., 2015).

*Mab-bsg* was identified from genome and transcriptome sequences. A fragment encompassing the full open reading frame was PCR amplified and used in a Gibson Assembly to generate a 3’ fusion with eGFP in a pSP expression vector (pSP-*Mab-bsg-eGFP*). RNA was *in vitro* transcribed using SP6 Polymerase (Roche), capping and polyA-tailing was performed using ScriptCap™ Cap 1 Capping System and Poly(A) Polymerase Tailing Kit (CellScript).

### Preparation of Lifeact-eGFP, Lifeact-mCherry, and Histone H1

Heterologous expression vectors for recombinant Lifeact-eGFP and Lifeact-mCherry were generated by cloning PCR-amplified constructs into pET-21a(+). The fragment encoding for Lifeact-eGFP was amplified from pT7-LifeAct-EGFP (Benton et al., 2013) using primer pair 5’-AAACATATGGGCGTGGCCGATCTGAT/5’-TTTTCTCGAGCTTGTACAGCTCGTCCATGC, digested with *Nde*I and *Xho*I, and cloned into pET-21a(+) to generate pET*-*Lifeact-eGFP. Similarly, a fragment encoding for mCherry was amplified from H2Av-mCherry (Krzic et al., 2012) using primer pair 5’-GAGGGGATCCTCGCCACCAGATCCATGGTGAGCAAGGGCGAGGAG/5’-GGTGCTCGAGGGCGCCGGTGGAGTGGCGGCC, digested with *Bam*HI and *Xho*I, and replaced the eGFP-encoding fragment in pET-Lifeact-eGFP to generate pET-Lifeact-mCherry.

Recombinant Lifeact-FP protein was expressed in *E.coli* BL21 after induction with IPTG (final concentration 1mM) at OD_600_=0.6-0.8. Cells were pelleted 4 hrs after induction, washed in PBS, and resuspended in lysis buffer on ice (50 mM NaPO4 pH 8.0, 0.5 M NaCl, 0.5% glycerol, 0.5% Tween-20, 10 mM imidazole, 1 mg/ml lysozyme). Resuspended cells were sonicated with 15-30 s pulses, centrifuged, and the supernatant mixed with equilibrated Ni-NTA agarose beads (Cube Biotech, Germany). Protein binding was carried out for 2 hrs at 4°C, beads were washed three times at high-salt/high-pH (50 mM NaPO4 pH 8.0, 250 mM NaCl, 0.05% Tween-20, 20 mM Imidazole), once at high-salt/low-pH (50 mM NaPO4 pH 6.0, 250 mM NaCl, 0.05% Tween-20, 20 mM Imidazole), and twice at high-salt/high-pH without detergent (50 mM NaPO4 pH 8.0, 250 mM NaCl, 20 mM Imidazole). Following the washes, beads were transferred into a poly-prep chromatography column (BioRad Laboratories) and the protein was eluted in multiple aliquots of elution buffer (50 mM NaPO4 pH 8.0, 150 mM NaCl, 250 mM Imidazole, 5% Glycerol). Collected protein fractions were analyzed by SDS-PAGE and dialyzed against PBS. Final concentrations were typically around 0.5 mg/ml; aliquots were stored at -80°C.

Histone H1 (Merck/Calbiochem) was fluorescently tagged using Texas Red™-X Protein Labeling Kit (ThermoFisher) as described (Mori et al., 2011). Final concentration was typically around 2 mg/ml; 10% saturated sucrose was added as anti-frost reagent, and aliquots were stored at -80°C.

### Immunohistochemistry

For whole mount *in situ* hybridization using NBT/BCIP as stain, embryos were heat fixed at indicated time points after developing at 25°C, devitellinized using a 1+1 mix of n-heptane and methanol, and post-fixed using 5% formaldehyde as described (Rafiqi et al., 2011a). For *in situ* hybridization using fluorescent Fast Red (Sigma-Aldrich), embryos were fixed with 4% formaldehyde and devitellinized using a 1+1 mix of n-heptane and methanol. For staining with Phallacidin and DAPI, embryos were fixed with 4% formaldehyde and devitellinized using a 1+1 mix of n-heptane and 90% ethanol. Whenever necessary, manual devitellinization was performed as described (Rafiqi et al., 2011a).

RNA probe synthesis, whole mount *in situ* hybridization, and detection was carried out as described (Lemke and Schmidt-Ott, 2009). The following cDNA fragments were used as probes: *Mab-ddc* (Rafiqi et al., 2010), *Mab-egr* (Kwan et al., 2016), and the newly cloned *Mab-Mmp1* and *Mab-bsg* cDNA fragments. Phallacidin staining was performed as described (Panfilio and Roth, 2010) with modifications: the stock (200 units/ml, Invitrogen B607) was diluted in PBS (1:25), embryos were stained for 3 hrs at room temperature and then briefly rinsed three times in PBS. DNA was stained using 4’,6-diamidino-2-phenylindole (DAPI, Life Technology D1306) at a final concentration of 0.2 µg/ml.

### Injections

Embryos were collected, prepared for injection, and injected essentially as described (Rafiqi et al., 2011b). dsRNA was injected with concentrations of 3.9 mg/ml, which corresponded to about 6 µM of *Mab-Mmp1* dsRNA. Concentration of injected *Mab-bsg* mRNA was about 3.3 mg/ml; concentration of injected Lifeact-mCherry protein was about 0.5 mg/ml, Histone H1 was injected at concentrations of about 0.7 mg/ml. Cas9 mRNA and all three sgRNAs were co-injected as a mix with a final concentration of 1 mg/ml of Cas9 mRNA and 50 ng/ml for each of the sgRNAs (Bassett et al., 2013).

### Microscopy

Embryos were embedded in a 3+1 mix of glycerol and PBS. Histochemical staining was recorded with DIC on a Zeiss Axio Imager M1 using 10x (dry, 10x/0.45); fluorescent staining was recorded by single-photon confocal imaging on a Leica system (SP8) using a 20x immersol objective (HC PL APO CS2 20x/0.75). Image stacks were acquired with a voxel size of 0.57 x 0.57 x 0.57 µm by oversampling in z.

### Confocal live imaging

Embryos were injected in the syncytial blastoderm stage with either recombinant Lifeact-mCherry (wildtype analyses) or a 1:1 mix of Lifeact-mCherry and *Mab-Mmp1* dsRNA. To restrict *Mab-bsg-eGFP* expression to the yolk sac membrane, *Mab-bsg-eGFP* mRNA was injected later and after germband extension had started. Time lapse recordings were started about 90-110 min after onset of germband extension for analysis of early, pre-disjunction tissue interaction, and 140-190 min after onset of germband extension for analysis of later, post-disjunction tissue interaction. Recordings were made by single-photon confocal imaging on a Leica system (SP8) using a 63x immersol objective (HC PL APO 63x/1.30 Glyc CORR CS2). Volumes were recorded in 15 to 20 confocal sections of 1 µm with simultaneous detection of mCherry and eGFP. Voxel size was 0.24 x 0.24 x 1 µm and volumes were collected at 20-second intervals for 10 min.

### Light-sheet microscope setup and imaging

Time-lapse recordings were performed using two Multiview light-sheet microscopes (MuVi-SPIM) (Krzic et al., 2012) with confocal line detection (de Medeiros et al., 2015). The microscopes were equipped with two 25 × 1.1 NA water immersion objective lenses (CFI75 Apo LWD 25XW, Nikon) or two 16 × 0.8 NA water immersion objective lenses (CFI75 Achro LWD 16XW, Nikon) for detection. Illumination was performed via two 10 × 0.3 NA water immersion objective lenses (CFI Plan Fluor 10XW). All objectives were used with the corresponding 200 mm tube lenses from Nikon. Fluorescence of mCherry was excited at 561 nm or 594 nm, TexasRed at 642 nm. Fluorescence was imaged simultaneously onto two sCMOS cameras (Hamamatsu Flash 4 V2) after passing corresponding fluorescence filters on the detection paths (561 nm LP, 647 nm LP, 594 nm LP, EdgeBasic product line, Semrock).

*M. abdita* embryos were injected in oil (refractive index 1.335, Cargille Labs) and mounted in an oil-filled fluorinate ethylene propylene (FEP) tube (Kaufmann et al., 2012). This tube was stabilized with a glass capillary that was placed into the capillary holder of the microscope. All embryos were imaged from four sides (one 90° rotation) every 1.5 or 2 min with a z-spacing of 1 μm for membrane labeled embryos and 2 μm for nuclear labeled embryos. The four orthogonal views facilitated a more uniform sampling, and the typical exposure time per plane of around 40 ms guaranteed an overall high temporal resolution. The resultant 4 stacks per time point were fused using previously published software (Krzic et al., 2012; Preibisch et al., 2010). All further processing and analysis was performed on the fused data sets. Analysis of *M. abdita* embryonic development was based on a total of 3 wildtype MuVi-SPIM recordings.

### Generation of embryo point clouds at and below the surface level

To allow for rapid image operations, fused 3D image stacks of individual MuVi-SPIM time points were transformed into point clouds. For this, fused 3D image stacks were read into Matlab using *StackReader*. Time-adaptive intensity thresholding was then used to segment the 3D image stacks into exactly two solid components: embryo and background. If segmentation returned more than one object, all but the largest one were eliminated and holes resulting from a lower fluorescence intensity in the yolk area were filled (*Segmentation*). To reveal fluorescent signal in layers below the embryo surface, the outermost layer of the segmented embryo was eroded using morphological operators *(imerode*) and a kernel radius of the specified depth. When needed, the surface was smoothened through morphological closing or opening (*imopen, imclose*). To visualize fluorescent signal for a specific layer of the embryo, the embryo was eroded at different depths and the resulting images subtracted from the original producing a set of concentric layers (*OnionCheat*). Point clouds were generated by mapping the geometrical voxel information of the segmentation (width, height, and depth) into vectors representing the 3D-cartesian coordinates [X,Y,Z], and their respective intensities into an additional vector (*PCBuilder*).

### Projections

To quantify tissue spreading over the full surface of the fly embryo, we used cylindrical projections as described (Krzic et al., 2012; Rauzi et al., 2015). Briefly, the anterior-to-posterior axis (AP) of the egg was calculated (*CovMat3D*) and aligned along the Z axis of the coordinate system (*VecAlign*). The cartesian coordinates were then transformed into cylindrical coordinates [X,Y,Z]->[θ,r,Z] (*Cart2Cyl*). For each position along Z and from 0 to 2π along θ, the mean intensities of all points between r_max_ and r_min_ were projected as pixels along width and height [W,H] of a two-dimensional image I (*CylProjector*). Translations and rotations (*PCRotator*) in the cartesian point cloud were used to position biological landmarks (e.g. dorsal/ventral midline) in I. To allow for a mapping of information obtained in I (tissue areas and cell tracks) back to the point clouds and stacks, the index information of all projected points was also stored in a vector array. Our cylindrical projection provided an approximate area conservation in the central domain of the embryo that was sufficient for visualization purposes. For quantitative analyses of serosa expansion, distortions were corrected at poles and furrows by using the law of cosines to weight the area of each pixel in I according to its contribution to the corresponding surface voxel in the embryo.

### Membrane segmentation

To quantify main aspects of cell shapes in fixed tissue, Phallacidin stained cells were segmented semi-automatically using Ilastik (Linux version 1.2.0) Pixel Classification framework (Sommer et al., 2011). In the case of mis-segmentation by the automatic algorithm due to lower resolution, cell outlines were corrected manually. Predictions were exported as a binary image stack. The spatial position of each cell within the imaged volume was defined by the centroid of the segmented cell. Individual cell volumes were extracted as a single connected component, the resulting objects were loaded as point clouds into Matlab and remaining holes were closed using *fillholes3d* with a maximal gap of 20 pixels (iso2mesh toolbox). To account for possible artifacts in image processing, objects smaller than 200 pixels and larger than 10000 pixels were excluded from further analyses. To reveal changes in cell and tissue dynamics in time-lapse recordings, individual cells and the expanding serosa were outlined manually.

### Feature extraction and quantification

Cell height was measured as object length orthogonal to the embryo surface. Cell surface area and cell circularity were measured by a 2D footprint that was obtained through a projection of the segmented cell body along the normal axis of the embryo. Cell tracks were obtained manually by following cells in selected layers of cylindrical projections. Germband extension was measured in mid-sagittal sections of time-lapse recordings: the most anterior point of the dorsally extending germband was used as reference, and germband extension was measured in percent egg length relative to the anterior-posterior length of the embryo.

### Cross correlation

A quantification of substrate and membrane movements was obtained using Matlab’s Computer Vision System Toolbox implementation of the Lucas-Kanade optical flow method on the respective layer projections. The orthogonal component corresponding to the AP axis was analysed on manually generated single cell segmentations, evaluating the area around the cell outline for the fluorescent signal of Lifeact-mCherry, and the entire cell area for the fluorescent signal of *Mab-bsg-eGFP*. The mean of the magnitudes was calculated for each frame and the cross correlation of both resulting vectors was evaluated for all 200-second time windows as described using Matlab’s Signal Processing Toolbox cross-correlation function (Goodwin et al., 2016). Negative controls were realized by calculating correlation between randomized cell and substrate area.

### Statistics

Statistical comparisons of correlation coefficients distribution were performed via Matlab implemented student’s t-test (two sided, unpaired) and hample’s test. P-values of tests are indicated in the figure legend.

### General image processing

3D reconstruction images of individual cells from z-stack segmentation data were done in Matlab (R2016b), images and stacks were processed using Fiji (2.0.0-rc-34/1.50a) and Matlab, and panels were assembled into figures in Adobe Photoshop and Adobe Illustrator. Custom Matlab functions for SPIM data processing are indicated with capital first letter and are available via github (https://github.com/lemkelab/SPIMaging).

## ACKNOWLEDGEMENTS

We thank A Guse and N Bloch for help with establishing a protocol for recombinant Lifeact-GFP; P Lenart for suggesting the use of labeled H1 and mRNA polyA-tailing; K Goodwin and G Tanentzapf for helpful suggestions on the cross-correlation analyses; M Benton, L Centanin, N Gorfinkiel, A Guse, K Panfilio, U Schmidt-Ott, J Wittbrodt, and members of the Lemke lab for discussions and/or comments on the manuscript. We acknowledge K Panfilio and two anonymous reviewers for their constructive feedback; I Lohmann, J Lohmann and J Wittbrodt for sharing laboratory equipment, and A Maizel for support. We are grateful to J Wittbrodt for his continuous and generous support. Funded by DFG grant LE 2787/1-1, HFSP grant RGY0082/2015, and a pre-doctoral HBIGS fellowship to LS.

## AUTHOR CONTRIBUTIONS

Conceived and designed experiments: FC, VN, EG, SL. Established SPIM imaging in *M.abdita*: FC. Performed selective plane microscopy: FC, DK. Build SPIM: DK, LH. Developed and implemented routines for SPIM data processing: EG. Evaluated SPIM data: FC, PG, EG, DK, LH, SL. Performed confocal microscopy: VN. Evaluated confocal data: PG, EG, VN, SL. Performed cross correlation analyses: EG. Performed molecular biology and histochemistry: FC, VN, MW. Provided data processing and IT infrastructure: EG, LS. Assembled figures and drew sketches: FC, PG, VN, SL. Drafted or wrote manuscript: FC, VN, SL. Read and commented on manuscript: all authors.

---

**Figure 1 supplement 1.**
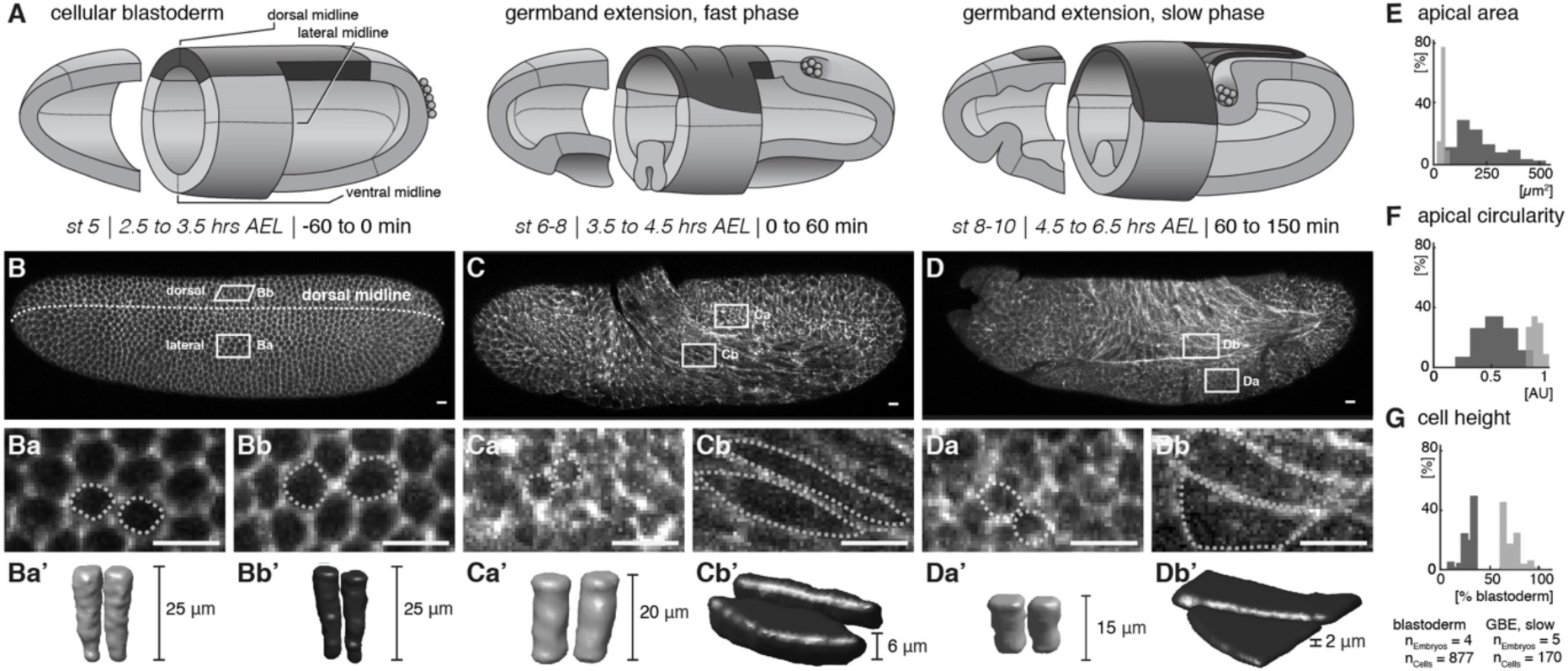
Quantitative analyses of cell measures in *M. abdita* embryos fixed at subsequent stages of development. (**A**) Model of early extraembryonic development in *M. abdita* without distinction of amnion and serosa (extraembryonic tissue labeled in black). (**B-Db’**) Global embryonic view of fixed embryos stained for Phallacidin to outline actin cytoskeleton (**B-D**), with close-up views (**Ba-Db**) and three dimensional volume renderings (**Ba’-Db’**) of embryonic (**Ba-a’, Ca-a’, Da-a’**) and extraembryonic cells (**Bb-b’, Cb-b’, Db-b’**). (**E-G**) Collective quantitative analysis of apical cell area (**E**), apical cell circularity (**F**) and relative cell height (**G**). Apical circularity (c) was defined as c=1 for a perfect circle and c < 1 for angular shapes with c = 4 π area/perimeter^2^ (Thomas and Wieschaus, 2004). Unless indicated otherwise, embryos and close-ups are shown with anterior left and dorsal up. Presumptive embryonic cells and corresponding quantifications are shaded grey, presumptive extraembryonic cells and corresponding quantification are in black. Scale bars, 10 µm.

**Figure 2 supplement 1.**
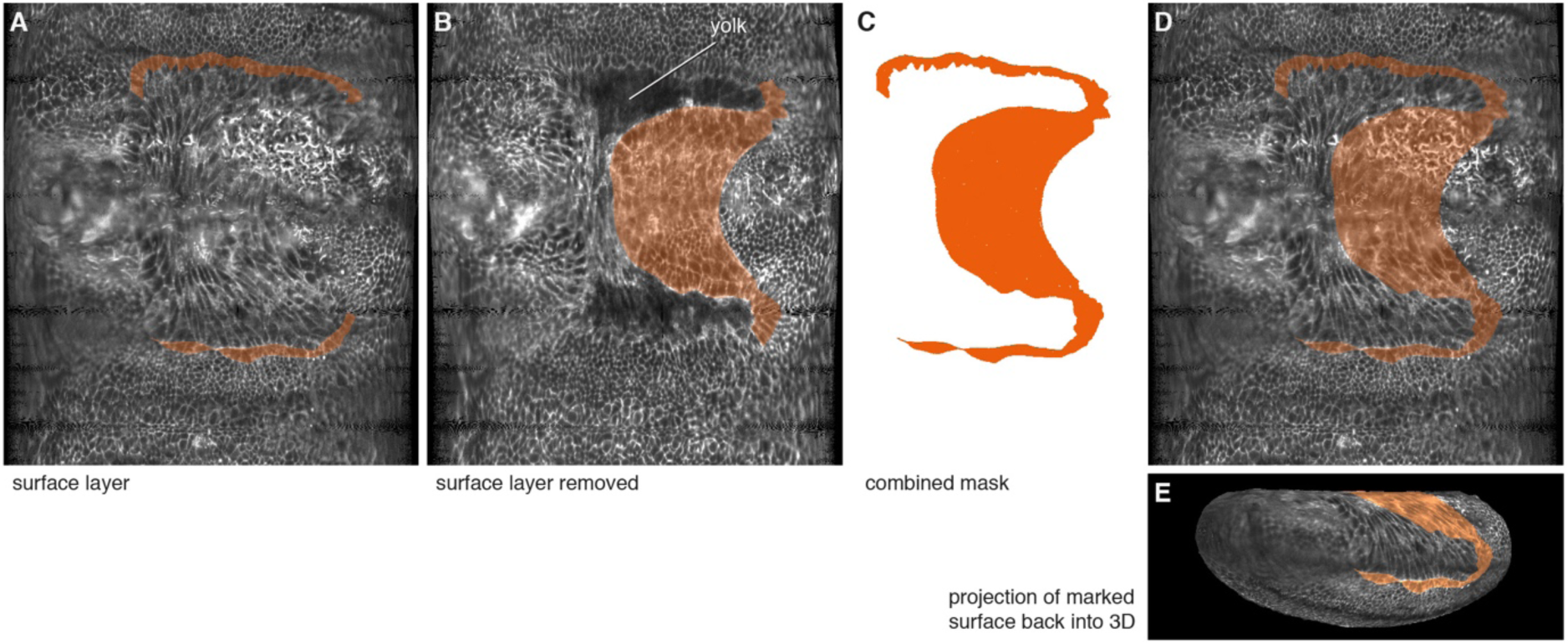
Surface layers were successively peeled off to mark the amnion. (**A**) Lateral amnion cells (one-to-two cells wide) were marked on the surface layer on an unrolled embryo (see material and methods). (**B**) The ventral amnion was marked on the underlying layer by computationally removing the surface layer. (**C**) Both masks were subsequently combined. (**D,E**) The combined mask was then mapped back onto the surface layer of the unrolled embryo (**D**), and the projection of the marked surface was mapped back into 3D (**E**).

**Figure 3 supplement 1.**
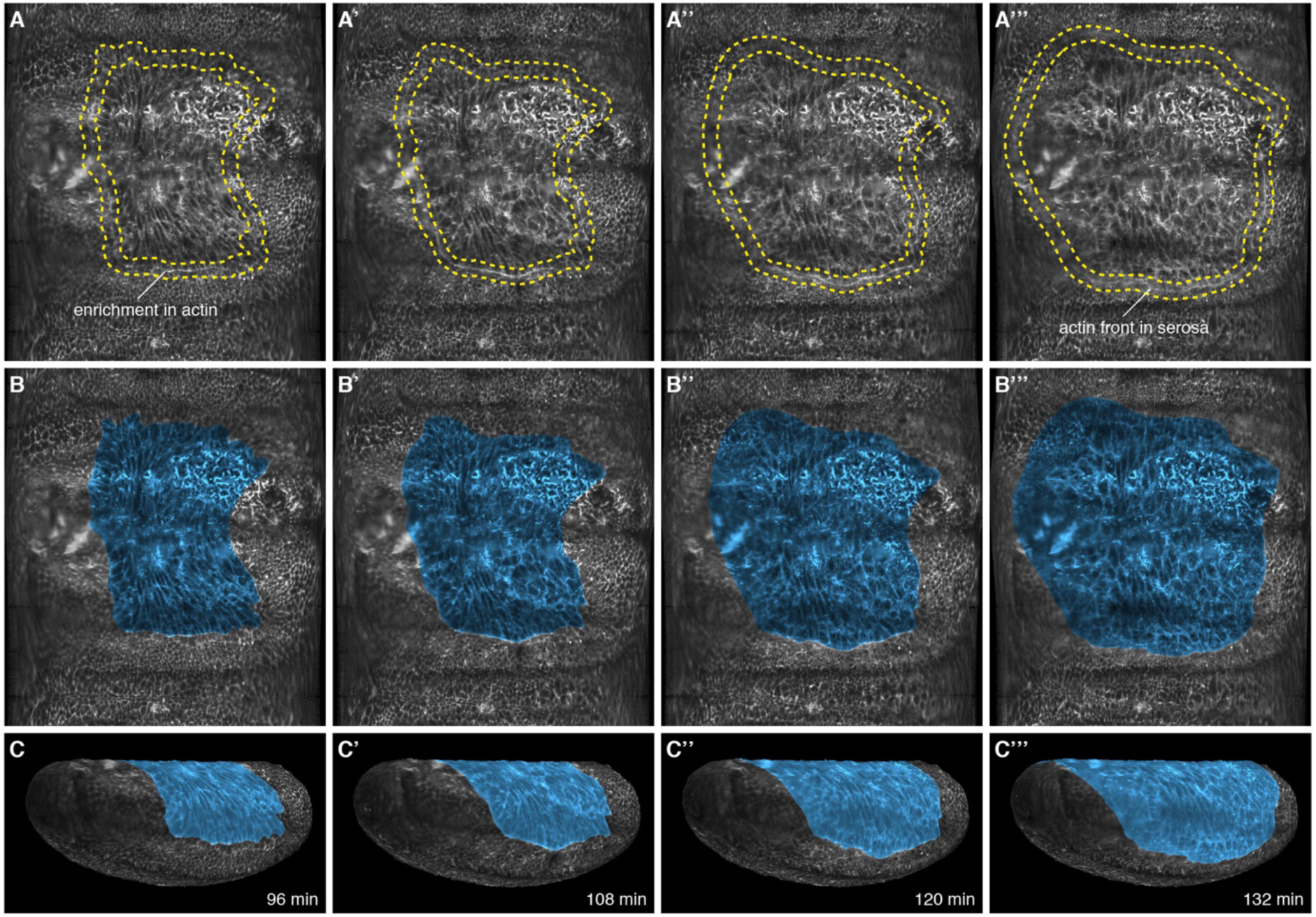
Cell tracks, cell shape, and enrichment of the actin reporter at the serosa periphery were used as landmarks to mark the amnion. (**A,B**) Marked outline (**A-A’’’**) and tinted area (**B-B’’’**) of serosa in 2D projected embryo surfaces. (**C-C’’’**) Projection of the marked surfaces into 3D renderings. Serosa disjunction was determined kinematically by visually following actin enrichment between serosa and amnion. As the actin cable moved towards the ventral side of the egg and continued migrating in front of the large serosa cells, the smaller cells of the presumptive amnion stopped moving. This process was best resolved by repeatedly playing recordings forward and backwards in time.

**Figure 4 supplement 1.**
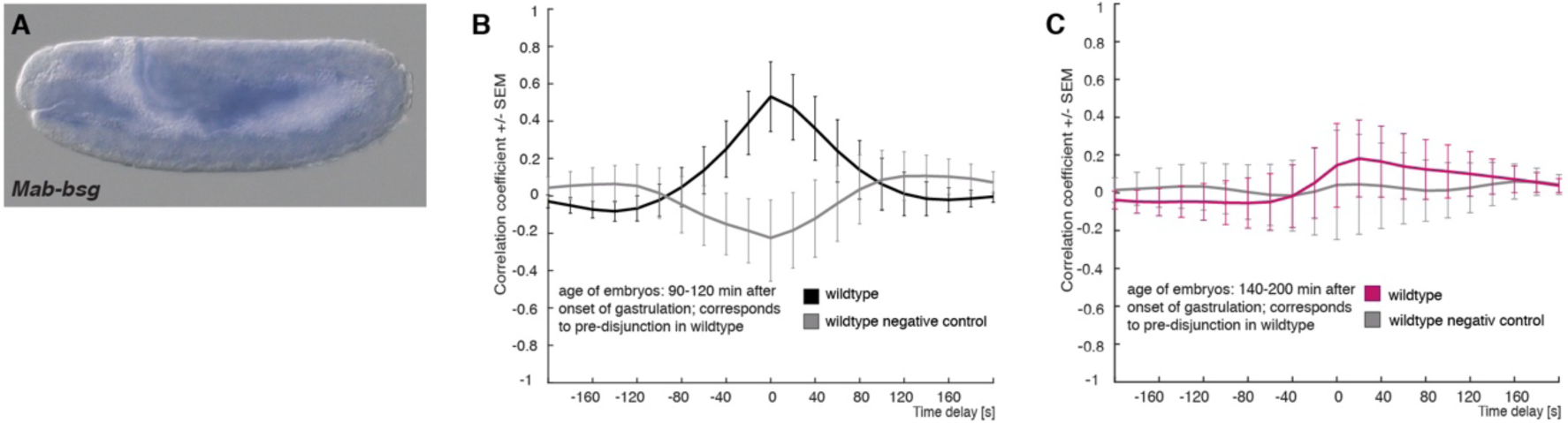
Expression of *M. abdita basigin* (*Mab-bsg*) and controls for cross correlation analysis in wildtype embryos. (**A**) Expression of *Mab-bsg* during germband extension stage. (**B,C**) Average cross-correlation function of corresponding cell and substrate movements in wildtype embryos as shown in Figure 4 for pre-disjunction (**B**, black) and post-disjunction (**C**, magenta); negative controls obtained by random pairings of cell and substrate movements from the same embryo revealed no detectable correlation (grey). Standard error of mean shown as bars (negative control 90-120 min: n = 20 cells from 5 embryos; negative control 140-200 min: n = 16 cells from 4 embryos).

**Figure 5 supplement 1.**
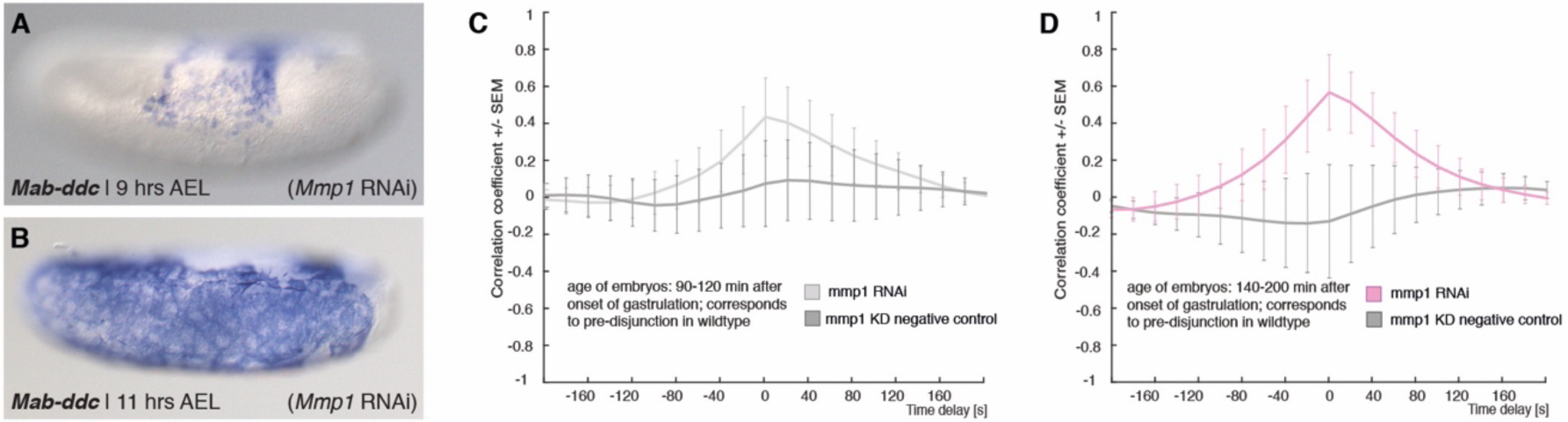
Expression of *Mab-ddc* as serosa marker and controls for cross-correlation analyses in *Mab-Mmp1* RNAi embryos. (**A-B**) Expression of *Mab-ddc* as serosa marker in embryos fixed 9 hrs AEL and 11 hrs AEL. (**C,D**) Average cross-correlation function of corresponding cell and substrate movements in *Mab-Mmp1* RNAi embryos as shown in Figure 5 for equivalents of pre-disjunction (**C**, light grey) and post-disjunction stages (**D**, light magenta); negative controls obtained by random pairings of cell and substrate movements from the same embryo revealed no detectable correlation (dark grey). Standard error of mean shown as bars (*Mmp1* RNAi negative control 90-120 min: n = 17 cells from 4 embryos; *Mmp1* RNAi negative control 140-200 min: n = 26 cells from 5 embryos).

**Figure 2 supplement, Movie #1**; movie corresponds to panels 2H-H’’. The *M. abdita* serosa folds over the putative adjacent amnion and then separates from it, as visualized in a SPIM time-lapse recording using Lifeact-mCherry as fluorescent actin reporter. Upper panel: “3D donut” cut (as indicated in Fig 2G); lower panel: corresponding 2D projection of surface view. Optical section (white line) and enrichment of actin reporter at serosa leading edge (in between yellow line) are indicated in the first frame. As the serosa expands over the putative amnion (117 - 135 min), the amnion appears to bend underneath the serosa. The serosa then separates from the adjacent tissue and continues to spread over the embryo proper. Time is relative to onset of gastrulation (0 min).

**Figure 2 supplement, Movie #2**; movie corresponds to panels 2I-I’’’. The *M. abdita* amnion remains lateral throughout germband extension and starts to progress towards the dorsal midline after onset of germband retraction, as visualized in a SPIM time-lapse recording using Lifeact-mCherry as fluorescent actin reporter. Upper panel: “3D donut” cut (as indicated in Fig 2G); lower panel: corresponding 2D projection of surface view. Optical section (white line) and putative amnion (in between orange line) are indicated in the first frame. As the germband retracts, actin protrusions are formed by the putative amnion cells (339-355 min). Time is relative to onset of gastrulation (0 min).

**Figure 3 supplement, Movie #1**; movie corresponds to panels 3A-C. *M. abdita* serosa spreading is a discontinuous process that is interrupted by a pause in tissue expansion, as visualized in SPIM time-lapse recording using Lifeact-mCherry as fluorescent actin reporter. Following manual marking of the serosa in 2D projections of the embryo surface (blue), surface and markings were projected back as 3D point cloud in lateral view (Figure 3 supplement 1). Serosa area was defined based on cell shape, cell tracking, and enrichment of the actin reporter in the periphery of the serosa. After onset of serosa spreading (30 min), a pause in area expansion becomes notable (60 - 110 min), while ventral closure is barely visible in the lateral view (190 - 236 min). Time is relative to onset of gastrulation (0 min).

**Figure 3 supplement, Movie #2**; movie corresponds to a ventral view of panels 3A-C. Ventral view of time-lapse recording and 3D projection as in movie #3. After passing of the poles (147 min), ventral closure can be clearly followed in the lateral view (190 - 236 min). Time is relative to onset of gastrulation (0 min).

**Figure 4 supplement, Movie #1**; movie corresponds to panel 4A and quantifications presented in 4E,F. Serosa cell and yolk sac movements prior to and after serosa disjunction. Dynamics of serosa cells (Lifeact-mCherry, magenta) and yolk sac (*Mab-Bsg*-eGFP, green) in wildtype embryos are shown before (left) and after (right) serosa disjunction. Each frame is an average z-projection of 4-7 confocal planes, which span the apical part of serosa cells and presumptive surface of the yolk sac membrane. Time lapse recording was performed with 3 frames/min for a total of 10 min; scale bar is 10 µm. Shown is a dorsal view with anterior to the left.

**Figure 5 supplement, Movie #1**; movie corresponds to quantifications presented in 5E-H. Serosa cell and yolk sac movements in *Mab-Mmp1* RNAi embryos. Dynamics of serosa cells (Lifeact-mCherry, magenta) and yolk sac (*Mab-Bsg*-eGFP, green) in *Mab-Mmp1* RNAi embryos are shown at stages corresponding to times before (left) and after (right) serosa disjunction in wildtype embryos. Each frame is an average z-projection of 4-7 confocal planes, which spans the apical part of serosa cells and presumptive surface of the yolk sac membrane. Time lapse recording was performed with 3 frames/min for a total of 10 min; scale bar is 10 µm. Shown is a dorsal view with anterior to the left.

